# A robust set of qPCR methods to evaluate adulteration in major spices and herbs

**DOI:** 10.1101/2024.01.11.575163

**Authors:** Marc Behr, Linda Garlant, Danilo Pietretti, Clément Pellegrin, Antoon Lievens, Ana Boix Sanfeliu, Alain Maquet, Lourdes Alvarellos

## Abstract

Spices and herbs are used in multiple forms by consumers and companies related to food or pharmaceuticals. Their intricate supply chains, high prices and limited productions make them susceptible to deceptive practises. However, robust methodology to detect multiple adulterants is still rarely available, preventing national control laboratories from enforcing European and national regulations.

This article presents qPCR methods developed to identify adulterants in six commonly consumed spices and herbs, namely paprika, turmeric, saffron, cumin, oregano and black pepper. Each genuine sample was blended with the top five adulterants identified in the European Union-wide coordinated control plan on herbs and spices and existing literature. These SYBR™ Green qPCR methods enable specific detection of adulterants, and their sensitivities allow to differentiate inadvertent contamination from deliberate adulteration. Methods performances were evaluated both in binary mixtures and with complex adulteration pattern, validating their effectiveness in safeguarding the authenticity of these high values commodities.

**Highlights:**

- Authenticity of spices and herbs is at high risk of fraudulent malpractices.
- We provide specific qPCR methods to detect prevalent adulterants.
- Trueness of each method was systematically assessed.
- Sensitivity of methods meet ISO requirements for purity assessment.
- Methods versatility is evaluated with multi-adulterated samples.

## 1. Introduction

Spices and herbs find extensive usage in food processing industry (75-80%), retail sector (15-25%) and the food service industry (5-10%) (Maquet et al., 2021). Beyond their culinary applications, these high-value products serve as essential ingredients in various food supplements and pharmaceuticals. This diversity of uses fuelled a rise on their global demand and put the supply chain at risk of malpractices and frauds, deceiving purchasers and consumers. The supply chains span across many countries, involving, from producers to consumers, a complex network of collectors, processors, traders, food manufacturers and retailers. Since malpractices might happen at any level of this intricate chain, analytical methods become imperative to ensure trust in the final product authenticity. Ensuring that marketed spices and herbs are “true-to-label” is not only essential for preventing consumer deception but also for certifying the absence of species causing food allergy or food intolerance, such as *Brassica juncea* and gluten-containing cereals. In such case, highly sensitive detection methods are crucial, as even minute amounts of allergens can trigger a severe allergenic reaction.

In absence of specific provisions for spices and herbs in the European Union (EU) regulatory framework, the assessment of authenticity and purity needs to be aligned with relevant ISO standards. To evaluate these ISO criteria, several analytical methods with complementary outputs are implemented, ranging from high-performance liquid chromatography coupled to mass spectrometry for non-authorised synthetic dyes and chemical markers (such as piperine for pepper, Sing et al., 2022; ISO 11027:1993) to X-ray fluorescence spectroscopy for elements detection such as heavy metals (Feng et al., 2021) and DNA-based analyses to highlight the presence of undeclared species (Gao et al., 2023; Hoffman et al., 2021). Hence, several sophisticated frauds have been evidenced with multistep workflows (Garber et al., 2016; Ordoudi et al., 2017; Xu et al., 2023). Among DNA-based techniques, quantitative PCR remains the gold standard for accurate and sensitive detection of extraneous/foreign matter of biological origin. Several research articles report the use of this technique in a broad range of food matrices, including spices and herbs such as *Capsicum annuum* (Kang, 2018), *Crocus sativus* (Villa et al., 2017), *Curcuma longa* (Oh & Jang, 2020), *Origanum vulgare* (Lievens et al., 2023) and *Piper nigrum* (Sousa et al., 2019). The aim of this work is to expand the available molecular diagnostic toolbox to additional adulterants and matrices.

Fit-for-purpose qPCR-based methods have been developed to estimate the quantity of five commonly known adulterants in five spices, including paprika (*Capsicum annuum*), saffron (*Crocus sativus*), cumin (*Cuminum cyminum*), turmeric (*Curcuma longa*) and pepper (*Piper nigrum*), as well as in the culinary herb oregano (*Origanum vulgare*). Based on the EU-wide coordinated control plan establishing the prevalence of fraudulent practices in the marketing of herbs and spices (Maquet et al., 2021) as well as scientific literature and reports (for instance (Khilare et al., 2019; Marieschi et al., 2012; Raina et al., 2024), five frequently found adulterants were selected for each main species. These selected botanicals were spiked into authentic spice/herb to reliably mimic the contamination or adulteration of commercial products. Subsequently, qPCR methods were either developed or optimized and assessed in the detection of contaminants and adulterants in binary mixtures. The specificity of primer pairs was systematically verified, as well as the performance of these qPCR methods with respect to their sensitivity, trueness, and overall suitability for analysing samples with complex adulteration patterns.

## 2. Materials and methods

### 2.1. Reference plant material and sample preparation

Whenever possible, plant reference material was sourced from living outdoor collections or seed banks, such as the Meise Botanical Garden (Meise, Belgium). Other carefully identified fresh and dry plant materials were collected from commercial sources or from private gardens. Additionally, dried plant materials were provided by various institutions (Supplementary Table S1). Individual plants or fruits were used whenever possible.

Reference plant DNA obtained from the Kew DNA bank (The Royal Botanic Gardens, Kew, UK), the DNA Bank of the Botanic Garden and the Botanical Museum Berlin (Germany) were primarily used for specificity tests. All DNA samples, as well as underlying voucher specimens, are deposited at the Botanic Garden and Botanical Museum Berlin and are available either via the Global Genome Biodiversity Network (Droege et al., 2014) or the Global Biodiversity Information Facility. Plant material and DNA were provided under the Convention on Biological Diversity 1992 agreement.

For total DNA extraction, fresh plant tissues were thinly sliced (< 1 mm) and oven dried (model UM500, Memmert, Schwabach, Germany) at 75 °C for 2 hours or until dry. Plant materials were milled using a MM301 ball mill (Retsch, Haan, Germany) for 2 minutes at 30 Hz using either 10 mL grinding jars and 10 mm beads (for softer and pre-ground materials) or 20 mL grinding jars and 20 mm beads (for harder materials such as seeds). Different sample blends were prepared from single species, their mixing ratio (mass/mass percentage) was determined gravimetrically (PG503-S, Mettler Toledo, Columbus, OH) to a final mass of 150 mg for each single species and blends (Supplementary Table S2).

### 2.2. DNA extraction, quality control and qPCR

A unique total DNA extraction protocol was applied to all samples. Powdered plant material (150 mg) was aliquoted and extracted with 1.25 mL hexadecyltrimethylammonium bromide (CTAB) buffer (MC1411, Promega, Madison, WI) containing proteinase K (ref 25530049, Invitrogen, Waltham, MA) and RNAse A (RNA-03, Omega Bio-Tek, Norcross, GA) at 100 µg/mL final concentration. After 30 seconds of vortexing, sample were incubated at 65 °C for 1 hour (1400 rpm shaking), followed by centrifugation (20 min at 17000 g). The supernatant was transferred to a fresh tube and a cleaning step performed with chloroform:isoamyl alcohol (24:1). After 10 seconds of vortexing and centrifugation (10 min at 17000 g), the aqueous phase was transferred to a fresh tube. A total of 350 µL of this aqueous phase underwent automated extraction and cleaning using a Tecan Freedom EVO liquid handler with Promega Purefood protocol and reagents (cell lysis buffer, ReliaPrep™ resin, prepared wash buffer, isopropanol, 50% ethanol and Promega elution buffer). The total extracted and washed DNA was eluted in a final volume of 150 µL.

DNA concentration was determined using a Qubit 4 fluorometer with Broad Range chemistry. The DNA Integrity Number (DIN) was determined with a gDNA screentape and a 4200 TapeStation instrument (Agilent, Santa Clara, CA). Finally, DNA purity was measured with a NanoDrop Eight spectrophotometer (Thermo Fisher Scientific, Waltham, MA).

qPCR amplification was carried out in a 20 µL reaction volume using 10 µL PowerUp™ SYBR™ Green Master Mix (ref A25777, Thermo Fisher Scientific), 400 nM of each forward and reverse primer and 25 ng of extracted DNA as template. The PCR program was as follow: activation of heat-labile uracil-DNA glycosylase (50 °C, 2 min), hot start activation (95 °C, 2 min), followed by 40 or 45 (for assessment of primer specificity) cycles of denaturation (95 °C, 15 sec) and annealing/extension (60°C, 1 min). A melt curve analysis was performed at the end of each run. The reactions were performed either with a QuantStudio 7 qPCR system (Applied Biosystems, Waltham, MA) in 96-well plates or a Rotor-Gene Q (Qiagen, Hilden, Germany) in 4-tube strips.

### 2.3. Primer design and specificity

For the detection of each plant species, primer pairs not available in the literature were designed based on publicly available nucleotide sequences (Benson et al., 2018). Primers used in this study are provided in Supplementary Table S3. The specificity of the 31 primer pairs for their intended target was assessed in technical duplicate by qPCR using a collection of template DNAs originating from more than 70 plant species related to adulteration.

### 2.4. Determination of a common reference gene for qPCR

DNA extracted from spices and herbs may contain several inhibitors (such as phenolics, proteins or sugars) that can affect downstream applications. A reference gene, which should be amplified on all species if the DNA extract is of sufficient quality, is therefore necessary to normalize the target DNA relative quantification (Bustin et al., 2009). Nine primer pairs targeting conserved reference regions were evaluated on a first collection of high-quality DNA from 46 plant species by qPCR. Out of these nine primer pairs, three of them (tRNA-Ile, 18S rDNA and 5.8S rDNA) showed positive amplification in every species, but only the tRNA-Ile primer pair showed rather stable C_q_ values, ranging from 16 to 21 for all tested species. qPCR using 18S rDNA and 5.8S rDNA primer pairs resulted in very low C_q_ values (< 8) for some species and C_q_ values between 18 and 20 for other species. Consequently, the tRNA-Ile primer pair was further tested using template DNA from 27 additional plant species with similar results and was therefore selected for normalization (Supplementary Table S4).

### 2.5. qPCR method validation

Validating a qPCR method requires to assess its robustness, sensitivity, trueness, as well as performing an inhibition test on investigated matrices. The concept of robustness in a method refers to its ability to yield consistent results even when slight deviations in experimental conditions are deliberately introduced. This parameter was evaluated by modifying the volume of enzyme in the mastermix (- 10%); primer concentration (-30%), as well as changing the annealing temperature (+1 °C or –1 °C), resulting in eight different reaction set-ups. Four of these set-ups were run on a QuantStudio 7 qPCR system, while the remaining four were run on a Rotor-Gene Q. DNA standard solutions containing 60 copies of the adulterant target gene (calculated from C-values available in the Kew databank, Leitch et al., 2019) were prepared in background DNA at 5 ng/µL and analysed in technical triplicates. Each method, in all tested conditions, resulted in successful amplification, thereby qualitatively validating the robustness of all qPCR methods (Supplementary Table S5). The inhibition test consists in the qPCR amplification of a serial 1 to 4 dilutions (from undiluted to 1/256) of pure plant DNA. The C_q_ value corresponding to an undiluted sample (expected without inhibition), extrapolated from the standard curve obtained with the serial dilutions, was compared to the measured C_q_ value of the undiluted sample. The serial dilutions and undiluted sample were analysed in technical duplicates and repeated on a separated plate. The sensitivity was determined in a 2-step approach. First, serial 1 to 10 dilutions (from undiluted to 1/10000 in background DNA) of binary mixtures (at 5 ng/µL) consisting in 10 % (m/m) adulterant in background matrix were analysed in technical triplicate, alongside a no template control (NTC, background DNA at 5 ng/µL). The two dilutions spanning the detection limit (i.e. amplification in all replicates for the higher concentration and either 0, 1 or 2 positive amplification(s) for the lower concentration) were selected for the second step. These two concentrations were then tested in triplicates and on three consecutive days. The limit of detection (LOD) was defined as the smallest concentration giving only positive amplifications with a C_q_ value lower than the NTC (Bustin et al., 2009). Finally, the trueness test aimed to assess the ability of the method to quantify the true proportion of one adulterant in a background matrix. Each pure herb or spice sample was separately admixed with the five most common adulterants, resulting in binary mixtures. The trueness was determined by analysing these binary mixtures composed of 2%, 5%, 50% and 100% (m/m) of adulterant with the background matrix. Additionally, samples consisting of a blend of each herb/spice (50% m/m) and its five adulterants (each 10% m/m) were prepared (six blends containing six botanicals in total). The binary mixtures were prepared in duplicate, and two independent extractions were used. Each was run on technical triplicates on two separate days. Amplification was performed using primer pairs targeting both the adulterant and the tRNA-Ile reference gene. Three quantification methods, adapted from Kang & Tanaka (2018), were evaluated to choose the best performing approach. In the selected method, the calibration curve is plotted with values calculated as ΔC_q_ = C_q target_ – C_q ref_ where C_q target_ is the test C_q_ value of the adulterant gene and C_q ref_ is the test C_q_ value of tRNA-Ile. The percentage of adulterant in the binary mixture is then calculated as 10^[(ΔCq-b)/a]^ where a and b are the slope and the intercept of the binary mixtures standard curve, respectively.

## 3. Results and discussion

### 3.1. Primers specificity

Specificity tests were performed on representative DNA samples encompassing authentic herbs, spices, as well as their commonly known adulterants and closely phylogenetically related species. Amplicon length was kept under 200 bp to enhance amplification efficiency, except for primers detecting *Fagopyrum esculentum* (buckwheat) and *Helianthus annuus* (sunflower) (amplicon lengths of 248 and 289 bp, respectively). Most primer pairs successfully amplified their target species with C_q_ values comprised between 20 to 25 cycles. In contrast, DNA from other species exhibited amplification differences of at least 10 cycles (Supplementary Table S6). The primer pair designed to detect *Origanum vulgare* amplifies the species allowed to be labelled as oregano (ISO 7925:1999), including *Origanum vulgare, Origanum vulgare* L. subsp. *hirtum*, *Origanum onites* L. and *Origanum* × *intercedens* (all with C_q_ values between 21 and 25 cycles). However, it also amplifies, to a lesser extent (C_q_ ca. 33 cycles), *Origanum majorana* (marjoram) which should not be labelled and sold as oregano (ISO 7925:1999). The primer pair designed to detect *Crocus sativus* (saffron) specifically amplifies this spice (C_q_ ca. 25 cycles), as well as *C. cartwrightianus* which has been identified as the sole progenitor of saffron (Nemati et al., 2019). The primer pair was not able to amplify neither *C. vernus* nor *C. speciosus*. The primer pair aiming at detecting *Cuminum cyminum* (cumin) were found to be specific for this spice (C_q_ ca. 20 cycles), with slight cross-amplification detected with *Arachis hypogaea* (peanut) (C_q_ ca. 30 cycles). This cross-reactivity was considered acceptable. *Carum carvi* (caraway) and *Bunium persicum* (black cumin or black caraway) were both amplified at higher C_q_ values (34 and 32 cycles, respectively), allowing the discrimination between these two species, commonly referred to as cumin in some countries, and *Cuminum cyminum*. The first primer pair designed to detect *Curcuma longa* (turmeric) showed the same amplification pattern with *C. zanthorrhiza*, displaying similar C_q_ values and melting temperatures. Since *C. zanthorrhiza* might be used as a turmeric adulterant, a new primer set was tested. This second pair (CuLo-CHS1, designed on *CHALCONE SYNTHASE 1* gene) turned to be more specific to *C. longa* but *C. zanthorrhiza* was still cross-reacting (26 cycles with *C. longa* versus 32 with *C. zanthorrhiza*). *C. zedoaria*, another possible turmeric adulterant, was not amplified. The primer pair targeting *Piper nigrum* (black, white, green and red pepper) was specific for this spice, with C_q_ values comprised between 20 to 22 cycles, while closely related *P. longum* (long pepper), which cannot be labelled as pepper (ISO 959-1:1998), was not detected.

The primer pair Pun1-p2, used for paprika authentication, was used to detect the presence of *Capsicum* species in turmeric and black pepper. This primer pair amplified DNA from *Capsicum annuum* (multiple cultivars of sweet bell peppers and chili peppers), *C. frutescens*, *C. baccatum*, *C. chinense* and *C. pubescens* (all chili peppers), therefore enabling the detection of several adulterants. The primer pair was not similarly efficient for all *Capsicum* accessions, since a first, large subset showed C_q_ values of 22-24, but two amplified around 28 cycles. This shift is probably due to the amplification of two products in the first subset, while only a single melting temperature was monitored for cultivars amplifying later. A slight cross-amplification was detected with *Physalis peruviana* (goldenberry) (ca. 30 cycles).

The Cru770 primer pair, targeting *Brassica* species (Mbongolo Mbella et al., 2011), provided a similar amplification pattern in *Brassica juncea* (brown mustard), *B. napus* (rapeseed) and *B. oleracea* (cauliflower). Contrasting with these sought-after cross-amplifications, the primer pair targeting *Tagetes* species (marigold) amplified some undesired species. These primers designed in the *LYCOPENE ε-CYCLASE* (*LCYe*) promoter region of *Tagetes erecta* (Mexican marigold) (Zhang et al., 2019) amplified successfully both *T. erecta* and *T. patula* (French marigold). However, since *LCYe* is a well-conserved gene in land plants, forming part of the carotenoid biosynthesis pathway, significant amplification was recorded in other species, such as *Helianthus annuus* (ca. + 3 cycles as compared to *Tagetes* species), *Calendula officinalis* (pot marigold) (+ 5 cycles) or *Carum carvi* (+ 10 cycles). Importantly, despite this broader amplification, the primer pair did not amplify *Crocus* DNA, thus not hindering its use for confirming the presence of *Tagetes* in saffron samples. Considering the lack of availability of public genomic data for *Tagetes* species, *LCYe* was retained as the target gene used to detect this possible adulterant.

Primers pairs deemed sufficiently specific for detecting possible contamination or adulteration were used in the subsequent steps of method validation.

### 3.2. Inhibition

Spices and herbs are known to contain a multitude of water-soluble specialised metabolites (such as polyphenols), and often undergo drying or boiling processes during their preparation, which may degrade DNA. Furthermore, the matrix in which adulterants are embedded may influence the detection and quantification of these contaminants. Additionally, obtaining DNA adulterants may also present difficulties, such as contamination or low yield, for instance in tissues undergoing programmed cell death. In this context, the inhibition test assesses the suitability of DNA extracted from spices, herbs or an adulterant for amplification. Following the MIQE recommendation (Bustin et al., 2009), inhibition was evaluated by comparing the expected and the observed C_q_ values in serial dilutions of DNAs (Figure 1).

**Figure 1.**
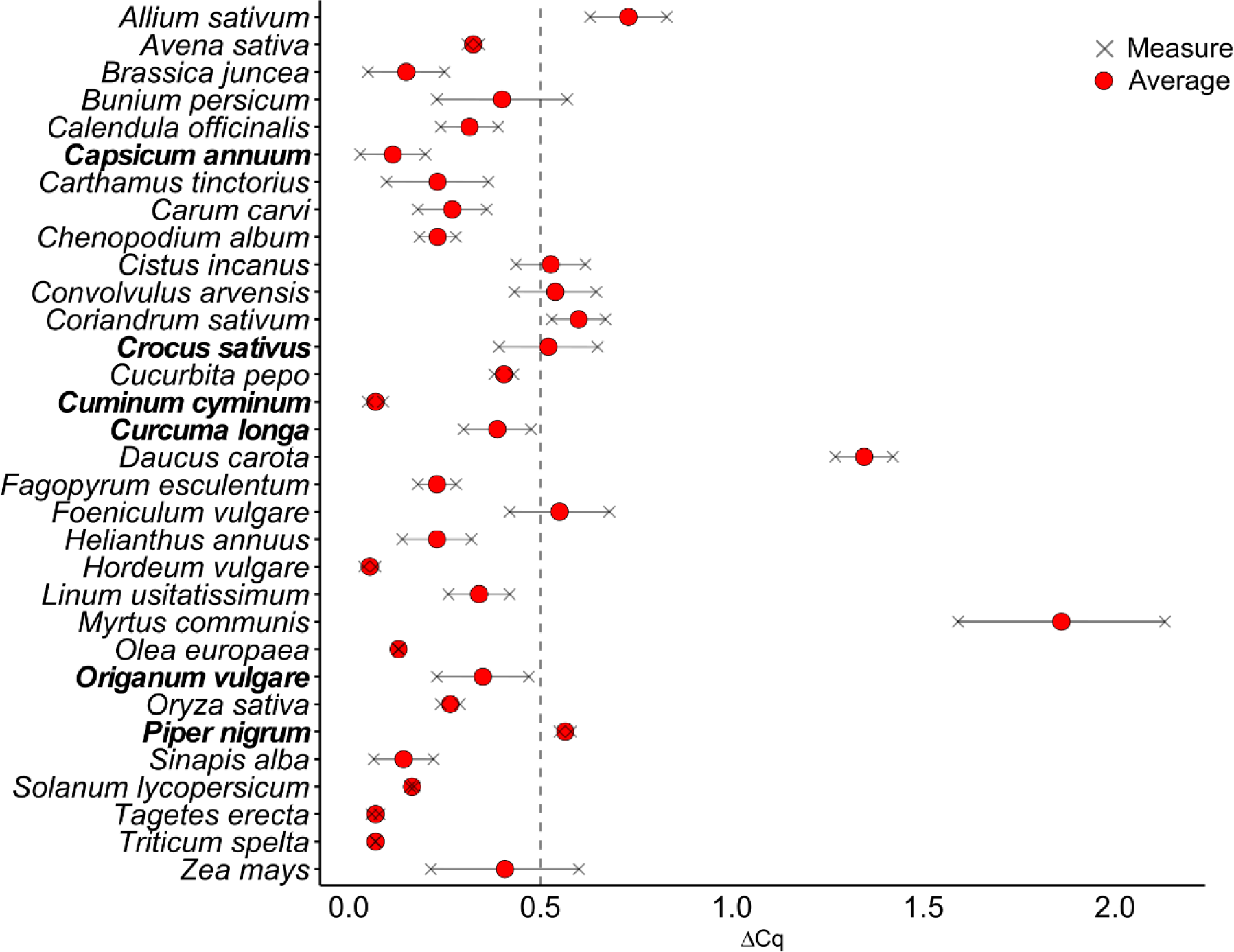
Inhibition test in herbs, spices and adulterants. ΔC_q_ consists in the difference between the expected and the observed C_q_ values in serial dilutions of DNA and should be lower than 0.5 (European Network of GMO Laboratories, 2015). Investigated spices and herbs are indicated in bold text. Data are provided in Supplementary Table S7.

DNA extracts from black pepper and saffron showed a risk of inhibition as their average ΔC_q_ is higher than 0.5. This finding could potentially affect the accurate quantification of adulterants present in these samples. Among the other botanicals, common myrtle (*Myrtus communis*), carrot (*Daucus carota*) and garlic (*Allium sativum*) exhibited a high potential for inhibition. *Cistus incanus*, field bindweed (*Convolvulus arvensis*), coriander (*Coriandrum sativum*) and fennel (*Foeniculum vulgare*) display risk of moderate inhibition (average ΔC_q_ close to 0.5). For samples containing a substantial proportion of these species, careful investigations are advisable. This can be achieved through preliminary DNA dilution before qPCR analysis or by normalizing the results using a reference gene, as further explained in paragraph 3.4. The potential for inhibition depends on the species but also on sample preparation and may also differ between plant materials from the same species. Therefore, the data reported in Figure 1 should not be considered as constant across all analyses but rather as indicative of the potential of inhibition of specific targeted adulterants.

### 3.3. Sensitivity

The sensitivity of primer pairs designed to detect potential adulterants in authentic spices and herbs was assessed in binary mixtures (authentic single species admixed with one corresponding adulterant). Sensitivity was quantified in terms of LOD observed in admixed mass samples (expressed as m/m percentage), fulfilling the applicability conditions (amplification performed in real matrices). The LOD determination consisted in two parts. The aim of the first part was to establish a mass concentration range (e.g. between 0.1% and 0.01% of adulterant in authentic material), for evaluating LOD in triplicates and over three days (second part). The results expressed as minimum percentage (m/m) of each detectable adulterant in the corresponding matrix are shown in Table 1.

**Table 1.** Sensitivity of qPCR methods for detecting foreign species in *Capsicum annuum*, *Crocus sativus*, *Cuminum cyminum*, *Curcuma longa*, *Origanum vulgare* and *Piper nigrum*. The percentages (m/m in authentic matrix) indicate the lowest detectable levels of foreign species in the nine replicates of the second part.

| ***Capsicum annuum*** |  | ***Crocus sativus*** |  | ***Cuminum cyminum*** |  |
| --- | --- | --- | --- | --- | --- |
| *Zea mays* | 0.01% | *Carthamus tinctorius* | 0.1% | *Carum carvi* | 0.01% |
| *Daucus carota* | 0.01% | *Curcuma longa* | 0.1% | *Coriandrum sativum* | 0.1% |
| *Solanum lycopersicum* | 0.001% | *Triticum spelta* | 0.1% | *Sinapis alba* | 0.1% |
| *Helianthus annuus* | 0.01% | *Tagetes erecta* | 0.01% | *Linum usitatissimum* | 0.01% |
| *Allium sativum* | 0.01% | *Calendula officinalis* | 0.01% | *Cucurbita pepo* | 0.1% |
| ***Curcuma longa*** |  | ***Origanum vulgare*** |  | ***Piper nigrum*** |  |
| *Zea mays* | 0.1% | *Olea europaea* | 1% | *Cuminum cyminum* | 0.01% |
| *Oryza sativa* | 0.1% | *Cistus incanus* | 1% | *Capsicum annuum* | 0.01% |
| *Triticum spelta* | 0.1% | *Chenopodium album* | 0.01% | *Oryza sativa* | 0.01% |
| *Avena sativa* | 0.1% | *Convolvulus arvensis* | 0.1% | *Allium sativum* | 0.01% |
| *Capsicum annuum* | 0.1% | *Myrtus communis* | 0.01% | *Brassica juncea* | 0.01% |

Most of the methods were able to detect foreign species within the mass concentration range 0.1%- 0.01%, with the highest accuracy achieved in the detection of tomato (*Solanum lycopersicum*) in paprika (0.001%). However, the methods performed poorly in detecting leaves from olive tree (*Olea europaea*) and cistus (*Cistus incanus)* in oregano samples, likely due to their low extraction yields compared to the oregano matrix and the presence of PCR inhibitors, as indicated by DNA quality controls (Supplementary Table S2).

The required purity level differs among the six investigated spices and herbs, necessitating careful consideration of the reported sensitivity data. These thresholds are defined in ISO standards specific for each species. ISO 7540:2020 reads that ground paprika may contain a maximal proportion of 1% extraneous matter, defined as species waste belonging to *Capsicum annuum* L. or *C. frutescens* L. plants. However, ISO 7540:2020 does not define a threshold for the quantity matter not belonging to *C. annuum* or *C. frutescens*, but instead provide a list of chemical requirements (such as the total capsaicinoids content) to which a paprika sample shall conform. Adulteration with other plant species can alter some of these chemical markers, making the qPCR pipelines described here an indirect method to detect contamination or adulteration. The sensitivities reported in Table 1, ranging from 0.01% m/m to 0.001% m/m, can detect the addition of unauthorised matter affecting the chemical specifications. In prior study, a qPCR method developed on garlic achieved a LOD of 0.001 ng/µL in serial water dilutions mixture containing equal quantities of DNA from chili pepper (*C. annuum*), garlic, onion, spring onion and ginger (Kang, 2018). This method aims at identifying the ingredients found in chili pepper powder and seasoned chili pepper sauces but was not developed to provide quantitative data.

ISO 3662-1:2011 stipulates that saffron of the highest grade (category 1) must contain less than 0.1% of foreign matter – defined as matter that is not belonging to *Crocus sativus*. The developed methods are able to detect the five investigated target species at this threshold (for safflower, turmeric and spelt) or at a lower content (0.01% for marigold and pot marigold). Using a primer pair designed on the *Internal Transcribed Spacer* region (*ITS*) of safflower (*Carthamus tinctorius*), similar results were previously obtained with qPCR on admixed saffron samples (Villa et al., 2017).

ISO 6465:2009 specifies that foreign matter in the three grades cumin - defined as the matter that is not part of *Cuminum cyminum* – should not exceed 0.5%. Notable, the five methods are able to detect the targeted species at either 0.1% (coriander, white mustard and pumpkin-related species) or 0.01% (caraway and flax). Interestingly, there are few methods describing DNA-based detection of adulterants or contaminants in cumin. Commercial RT-PCR tests were used to pinpoint the presence of peanut and hazelnut in cumin samples (Garber et al., 2016), while two qPCR assays were developed for the detection of *Prunus* species (Walker et al., 2018). More widely used methods include antibody-based tests, mass spectrometry or molecular spectroscopy (Cruz-Tirado et al., 2023; Garber et al., 2016). The five methods described in this article will therefore allow the detection of the adulterant target species at significantly lower concentrations as compared to previously reported techniques.

ISO 5562:1983 specifies that whole turmeric should contain less than 2% of extraneous matter – defined as substances not belonging to the *Curcuma longa* plant and part of plants other than the rhizomes of *Curcuma longa*. The targeted species (maize, rice, spelt, oat and bell pepper) were all detected at 0.1%. Previous investigations also reported a cut-off value of 0.1% of maize in turmeric powder by using qPCR with primer pairs targeting the *matK*, *atpF* and *ycf2* regions (Oh & Jang, 2020).

ISO 7925:1999 outlines that processed and semi-processed oregano should not contain more than 1% and 3% of extraneous matter, respectively – defined as matter not belonging to the leaves of oregano and all extraneous matter of animal, vegetable and mineral origin. While leaves from olive tree and *Cistus incanus* are detected at this maximal level, field bindweed (*Convolvulus arvensis*) on one hand, and goosefoot (*Chenopodium album*) and common myrtle on the other hand, are detected at 0.1% and 0.01%, respectively.

Finally, ISO 959-1:1998 specifies that processed black pepper should not contain more than 1.5% of extraneous matter – defined as all materials other than black pepper berries, regardless of whether they have vegetable or mineral origin. The five investigated species (cumin, chili pepper, rice, garlic and brown mustard) were detected at 0.01%. Using a different experimental setup, Sousa and colleagues (2019) were able to detect the presence of 0.0096% of maize (*trnL* region) and 0.0424% of *Capsicum* (*psbA*-*trnH* region) in black pepper by qPCR. These calculated contents derived from DNA standard solutions containing 2 and 10 haploid genome equivalents of maize and *Capsicum* in black pepper background DNA, respectively, representing the lowest values resulting in a positive amplification in all the technical replicates. However, such testing does not account for the differential DNA extractability of both the target and the background species in complex or binary mixtures. Nevertheless, both methods exhibit similar sensitivity for the detection of maize and chili pepper either as contaminants or adulterants.

### 3.4. Trueness

Analytical methods used in official tests should be capable to distinguish accepted contaminations (see ISO standards mentioned in paragraph 3.3.) and deliberate additions of matters not permitted in authentic spices and herbs. The uncertainty associated with a determined quantity of adulterant in an authentic product should not prevent to make such a distinction. Based on authentic material admixed with four quantities of adulterant (2%, 5%, 50% and 100%), the relative bias associated with each of these concentrations was evaluated by qPCR (Figure 2).

**Figure 2.**
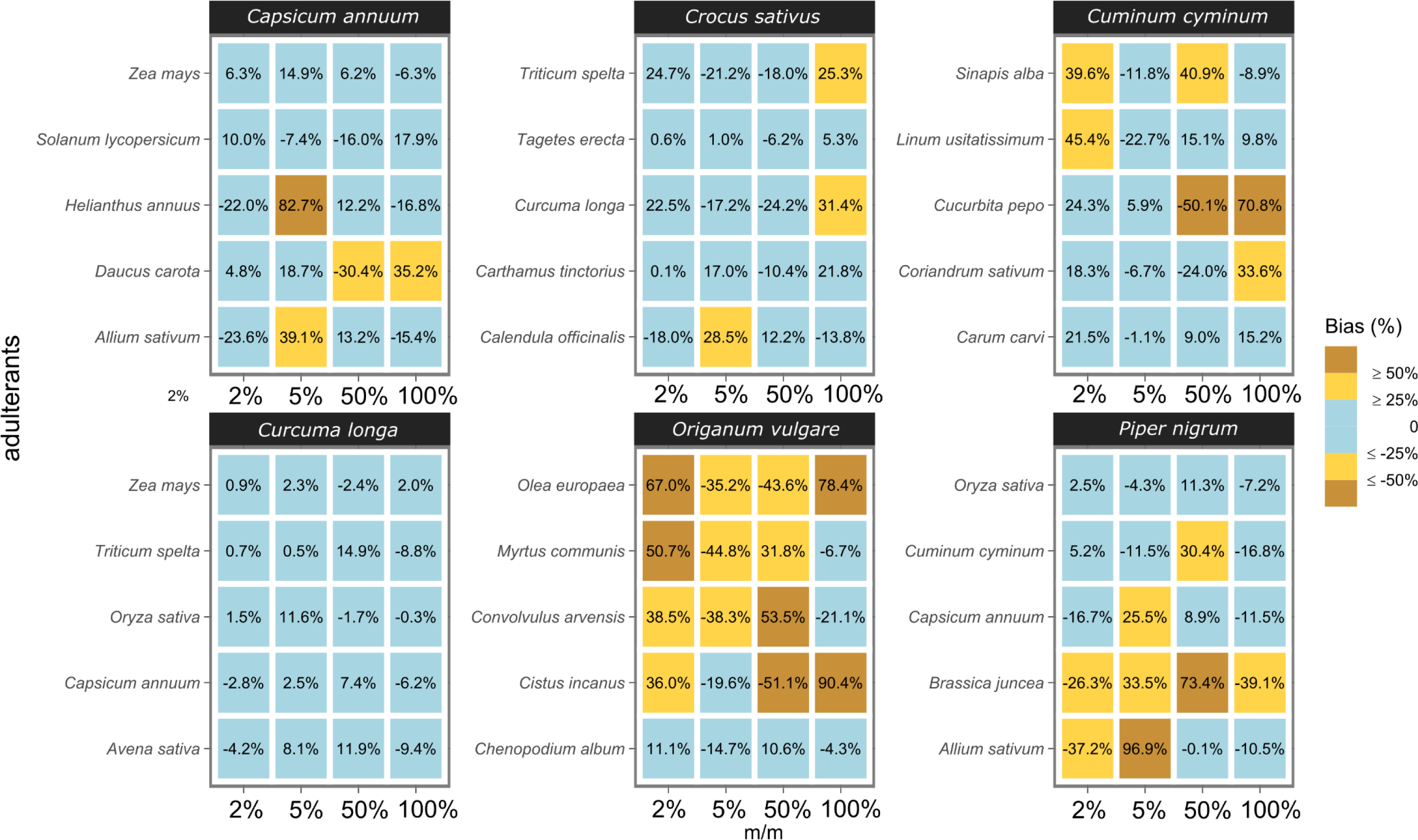
Relative bias associated to quantification of adulterants in paprika, saffron, cumin, turmeric, oregano and black pepper.

Most reported trueness values (86 out of 120) fall within ± 25% of the reference value, which is considered as an acceptable threshold in qPCR systems (European Network of GMO Laboratories, 2015). However not all methods perform equally well. For example, all the biases in turmeric are within the ± 25% interval, whereas only eight fulfil this condition in oregano. Oregano samples admixed with *Olea europaea* and *Cistus incanus* were not reliably quantifiable, likely due to the similar factors explaining the low sensitivity of these methods (low DNA extraction yield, presence of inhibitors). Smaller biases but still out of the acceptance criteria were achieved for field bindweed and common myrtle, while goosefoot met the acceptance criteria. These contrasting results within the same oregano matrix emphasize that precise quantification depends primarily on the composition of the adulterant (DNA quantity, integrity and extractability, presence of PCR inhibitors) rather than the matrix itself. Specifically, the main difference between goosefoot (meeting the acceptance criteria) and other oregano adulterants lies in the DNA extraction yield. The goosefoot DNA was repeatedly extracted with concentrations over 150 ng/µL while the DNA concentration ranged from 5 ng/µL for common myrtle to 30 ng/µL for field bindweed. Oregano DNA was quantified at 30-50 ng/µL, depending on the accession. Adulterants with low DNA yields are therefore more difficult to detect in the oregano matrix. Secondly, DNA preservation was better in goosefoot’s extract, with a DIN of 5.3, while field bindweed and olive tree displayed DINs of 3 and 2.1, respectively. Finally, no significant difference between the extracts were observed in the A_260_/A_280_ and A_260_/A_230_ ratios (Supplementary Table S2). The quantification bias of garlic in *Capsicum annuum* and *Piper nigrum* also exhibited values falling outside the acceptance criteria (Figure 2). As garlic yields more DNA than these two matrices (Supplementary Table S2), its DNA is dominant in these respective binary mixtures, implying that relative quantification by qPCR will be less accurate yet sensitive for low to very low amounts of garlic (2 and 5% m/m, Figure 2).

Normalisation with the tRNA-Ile reference gene, accounting for all the previously mentioned factors, improved the overall accuracy of the results, with few exceptions where the bias was lower without normalisation. In one specific case (oregano adulterated with common myrtle), estimating the proportion of myrtle was impossible without measuring the relative quantity of tRNA-Ile. Besides its poor yield (usually < 5ng/µL), extracted myrtle DNA displayed low A_260_/A_230_ ratio (0.79 and 0.84 in samples used for the trueness test), preventing accurate quantification in samples admixed with more than 50% of this plant material. The average C_q_ value of the target gene was 21.0 in 50% admixed samples, but 30.1 in the pure myrtle DNA. The C_q_ values of tRNA-Ile displayed the same trend, highlighting the low amplificability of samples containing high proportions of common myrtle leaves. Including a reference gene in the experimental procedure is, therefore, highly recommended, especially with plant materials that are difficult to extract and/or that contain high level of PCR inhibitors. Nanodrop absorbance ratio emphasized the presence of significant amounts of contaminants in many samples (Supplementary Table S2). One may expect improved precision with purer samples, provided that the modified preparation does not degrade the DNA (lower DIN scores) or lower the adulterant DNA relative content in the extract. The extraction protocol used in this article aimed at preparing large numbers of samples with a single semi-automated method. Depending on the matrix to be analysed, optimisation of DNA extraction method may lead to higher precision and sensitivity.

### 3.5. Detection of multiple adulterants in a single sample

The qPCR methods were tested for detection and the quantification of five adulterants simultaneously admixed in paprika, saffron, cumin, turmeric, oregano and black pepper matrices (Table 2).

**Table 2.**
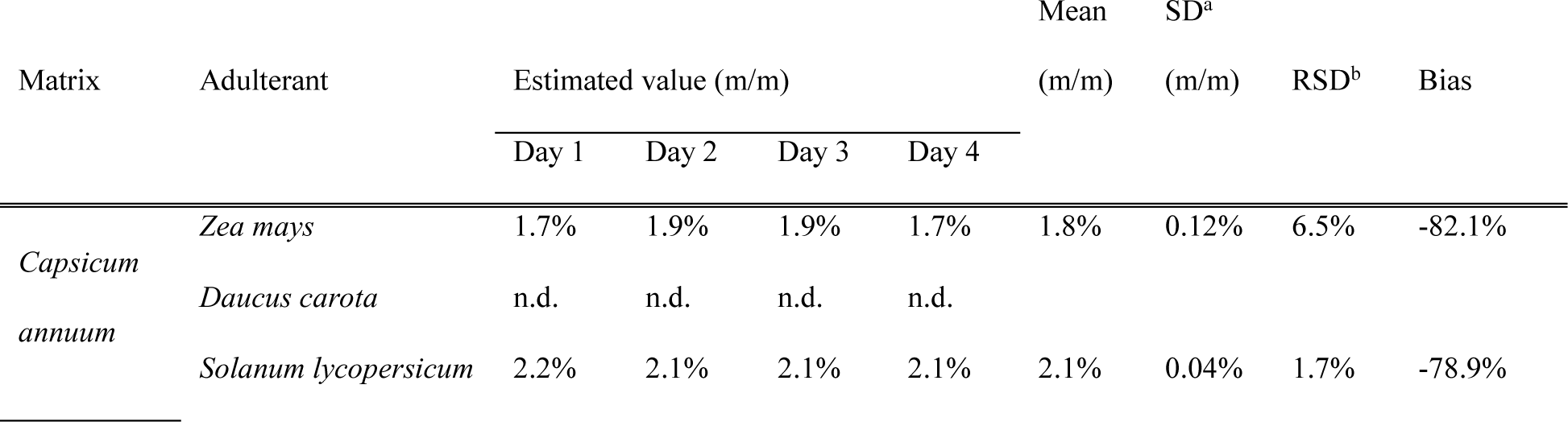
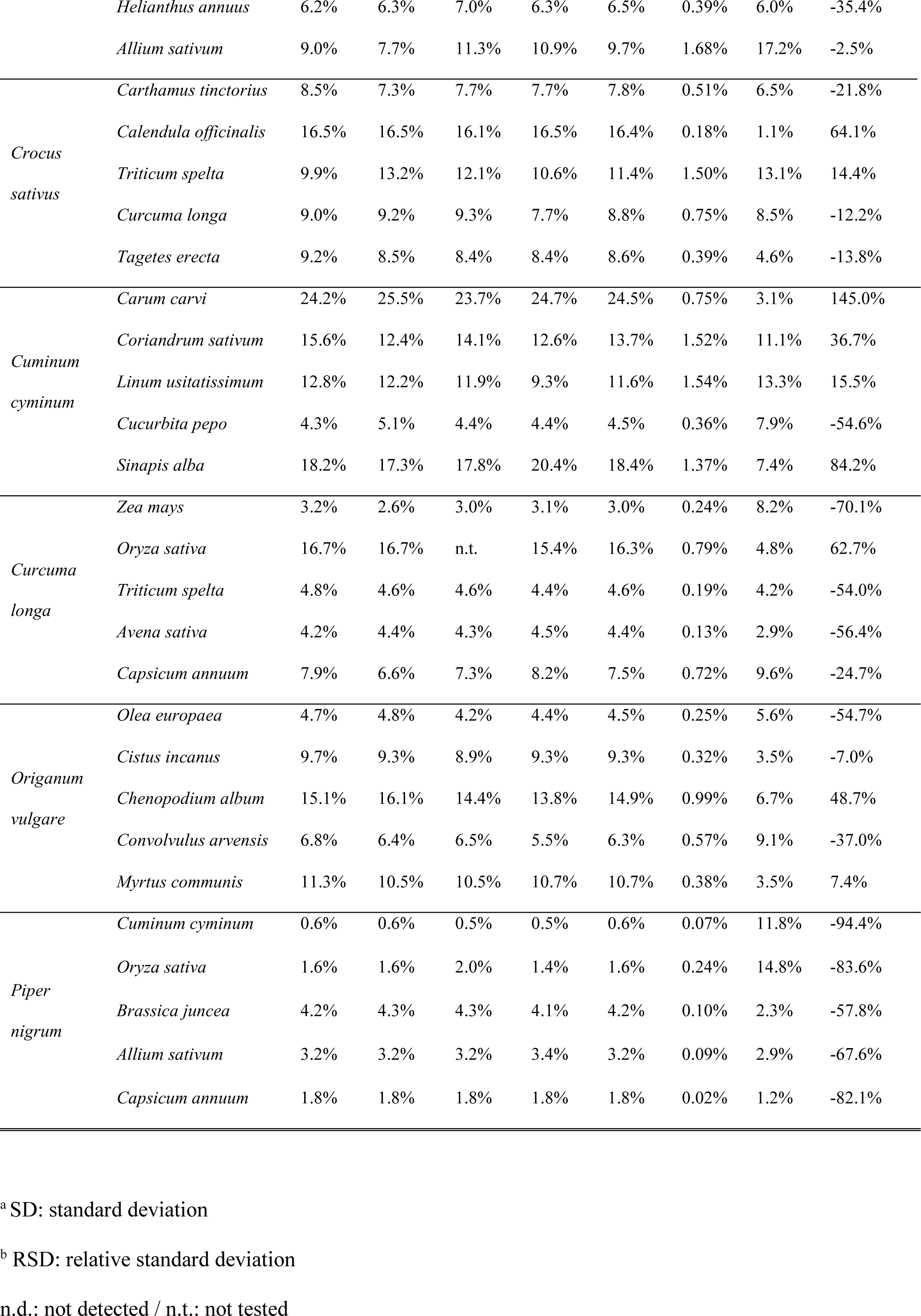
Quantitative analysis of complex herbs and spices admixed samples. Each matrix (50% m/m) was admixed with five main adulterants (each 10% m/m). Estimated values (based on target C_q_) were calculated using a daily calibration curve that consisted of binary mixtures with adulterants at 0.5%, 2%, 5%, 10%, 50% and 100% (m/m), repeated over four days. Each estimated value represents the average of technical triplicates.

With exception of carrot in paprika, all the targeted species were detected in the complex mixtures. The methods, therefore, demonstrate their capability to handle multi-adulterated samples. However, the bias associated with the quantification of these species is considerably higher than in binary mixtures. Several factors contribute to this discrepancy. First, there is no normalization with a reference gene, such as tRNA-Ile, as was done for the trueness assessment. Consequently, the estimated values remain uncorrected for the presence of inhibitors or DNA degradation. Second, the complexity of the mixtures leads to differential DNA extraction yields among the six species present in the sample. For instance, the absence of amplification of the target gene for carrot is likely due to its low DNA proportion in the extract. Third, the calibration curves used for estimating values were obtained with binary mixtures, which do not fully account for the complexity of the samples. Finally, unlike in trueness assessment, a single DNA extraction was performed for these samples; the precision of the estimation may, therefore, improve by conducting additional replicates. In conclusion, in the context of official analyses for fraud detection, quantification of complexed samples may be improved at a reasonable experimental cost by (1) normalising the target gene C_q_ with a reference gene such as tRNA-Ile and, (2) performing twice the DNA extraction from the matrix to provide an independent additional measurement.

## 4. Conclusion

The qPCR methods proposed in this study detect the presence and estimate the proportions of multiple adulterants in six commonly used spices and herbs. The applicability of these methods primarily relies on using botanicals binary mixtures for the assessment of LOD and trueness. The specificity of the primers and the potential for inhibition of each matrix and adulterant were meticulously investigated to highlight the methods limits and how to handle them. Normalisation with the reference gene tRNA-Ile resulted in improved precision and effectively mitigated issues arising from inhibitory matrices. Furthermore, all the methods rely on the cost-effective and widespread SYBR™ Green dye, providing an affordable and accessible approach for most laboratories. The approach of adulterant quantification developed here could be extended to other matrices and botanicals, thus supporting quality controls in the field of spices and herbs. Full internal validation and interlaboratory comparison will provide the measurement uncertainties associated with the methods, allowing their deployment for official analyses.

## Acknowledgements

The authors extend their gratitude to the Meise Botanical Garden (Meise, Belgium), The Royal Botanic Gardens (Kew, UK) and the DNA Bank of the Botanic Garden and Botanical Museum Berlin (Germany) for providing botanicals and DNA samples. They particularly want to thank Laure Ohleyer (Chambre d’Agriculture Côte-d’Or, France) for the generous donation of *Brassica juncea* seeds.

## Appendix A. Supplementary data

Supplementary data are available for this article.

